# DeLTa-Seq: direct-lysate targeted RNA-Seq from crude tissue lysate

**DOI:** 10.1101/2020.09.15.299180

**Authors:** Makoto Kashima, Mari Kamitani, Yasuyuki Nomura, Hiromi Hirata, Atsushi J. Nagano

## Abstract

Using current mRNA quantification methods such as RT-qPCR and RNA-Seq, it is very difficult to examine thousands of tissue samples due to cost and labor of RNA extraction and quantification steps. Here, we developed Direct-RT buffer in which homogenization of tissue samples and direct-lysate reverse transcription can be conducted without RNA purification. We showed that appreciate concentration of DTT prevented RNA degradation but not RT in the lysates of several plants’ tissues, yeast, and zebrafish larvae. Using the buffer, direct reverse transcription on the lysates could produce comparable amount of cDNA with that synthesized from purified RNA. Furthermore, we established DeLTa-Seq (**D**ir**e**ct-**L**ysate reverse transcription and **Ta**rgeted RNA-**Seq**) method. DeLTa-Seq is a cost-effective, high-throughput and highly-precise quantification method for the expressions of hundreds of genes. It enables us to conduct large-scale studies using thousands of samples such as chemical screening, field experiments and studies focusing on individual variability.

## Introduction

Quantification of gene expression is a popular approach to study various biological phenomena^1–8^. Recently, large-scale transcriptome analysis by RNA-Seq of hundreds of samples has been emerged in plant biology^9–12^. Although sequencing cost has been reduced drastically in the last decade^13^, the requirements for labor and cost at library preparation step obstruct routine use of RNA-Seq. To overcome these difficulties, several cost-effective RNA-Seq library preparation methods were developed^14–19^. Particularly, the two methods developed based on the 3’ RNA-Seq protocols for single cell RNA-Seq, Lasy-Seq^19^ and BRB-Seq^18^, enables us to early-pooling of samples resulting in reducing labor and cost into about two dollar per sample. In this situation, RNA extraction is remained as the most laborious and costly step in RNA-Seq. Especially, because plants tend to contain large amount of polysaccharide and high RNase activity, it costs more expensive to purify RNA from plant samples than animal cultured cells^20^. To obtain high quality RNA from plant samples, we homogenize and lyse the sample in the buffer containing guanidine hydrochloride to denature RNase, following by RNA purification with phenol chloroform extraction, silica column or magnetic beads that selectively bind to nucleic acid. If these RNA extraction steps can be skipped, the required cost and labor for RNA-Seq will be reduced more and make it easier to use RNA-Seq in large-scale studies with thousands of samples. Although several studies showed that RT can be possible directly from lysate of cultured animal cells in cell-lysis buffers^21–23^, in our knowledge, no study showed direct-lysate RT from crude tissue lysates. A previous study showed that reducing regents such as dithiothreitol (DTT) irreversibly deactivate intrinsic RNase in wheat^24^. Yet, whether DTT can portect RNA in the lysate of other speiseses and whether DTT does not inhibit reverse transcription has not been examined.

There is the tread-off between comprehensive gene quantification and sequencing cost^25^. For example, in the case of human transcriptome, it is estimated that ∼40 million reads are required for the reliable quantification of moderately-abundant transcripts, and as many as 500 million reads are required to comprehensively quantify rare transcripts such as transcription factors^26, 27^. In some cases, it is enough to focus on the limited number of genes of interest. When we focus on the limited number of genes, we can measure gene expression more accurately with small number of reads. This will enable us to examine biological phenomena with higher resolution of time, space, perturbation and individual variability. Thus, cost-effective methods of gene quantification such as targeted RNA-Seq and its relevant techniques to quantify hundreds of transcripts^29–32^ are required to understand complex behaviors of biological systems.

Here, we showed that high concentration of DTT can inhibit intrinsic RNase in plant and animal crude tissue lysates but not a reverse transcriptase. Using Direct-RT buffer containing DTT, we succeed in direct-lysate RT from plant, yeast and animal samples and RNA-Seq using the lysate. Finally, we combined direct-lysate RT and targeted RNA-Seq (DeLTa-Seq). DeLTa-Seq can drastically reduce cost and labor of RNA-Seq, and keep accuracy in quantification. DeLTa-Seq will pave the way for a large-scale research.

## Results

### Reducing regents inhibit RNA degradation in plant, yeast and animal lysates

For direct-lysate RT from plant tissues, we tried to apply the three regents [CL buffer^23^, CellAmp (TaKaRa, Kusatsu, Japan) and SuperPrep (TOYOBO, Osaka, Japan)] designed for direct-lysate RT from cultured mammalian cells. *Arabidopsis thaliana* seedlings were homogenized in these regents followed by an hour incubation at 22 °C. Then, RNA was purified from the lysates and checked with electrophoresis. Frozen rice (*Oryza sativa*) leaves were also homogenized with zirconia beads and dissolved in the regents. Incubation and RNA purification were conducted same as above. In both cases, any peaks of 18S and 28S rRNA were not observed, but short fragments of degraded RNAs were observed (Supplemental Fig. 1). These electropherograms indicated that these regents for direct-lysate RT from cultured mammalian cells cannot inhibit RNA degradation in plant lysates.

In the previous research^24^, inactivation of RNase in an extract prepared from dark grown wheat required 0.5 mM DTT or more than 0.014% 2-mercaptoethanol (2ME)^24^. Thus, we examined the ability of reducing reagents to protect RNA in plants’ lysates from RNase. We prepared 90 mM Tris-HCl (pH7.6) containing 10 mM, 50 mM, and 100 mM DTT and 2.5% 2ME, and homogenized *A. thaliana* seedlings in these buffers. After an hour incubation at 22 °C, RNA was purified from the lysates, followed by quality check with electrophoresis. In the electropherogram of the purified RNA from the lysate in negative control buffer (no reducing regents), no peaks of rRNAs were observed (Fig. 1), indicating that RNA was degraded in the lysate. In contrast, the clear peaks of rRNAs were observed in the electropherograms of the purified RNA from lysates containing DTT or 2ME. In the 10 mM DTT buffer, slight degradation of rRNA was observed in the region shorter than the 18S rRNA (Fig. 1). In the higher concentration of DTT or 2.5% 2ME buffer, degradation of RNA was inhibited (Fig. 1). In addition to *A. thaliana* seedling, we examined the effect of adding reducing regents to the lysates of *O. sativa* leaf and root and *Triticum aestivum* coleoptile with first leaf. Frozen rice leaves and wheat coleoptile were homogenized with zirconia beads, and Tris-HCl buffers containing the reducing regents were added. Like the results of *A. thaliana,* after an hour incubation at 22 °C, degradation of rRNA was observed in rRNA purified from the lysates in the buffer without any reducing regents and 10 mM DTT but not 50 mM and 100 mM DTT and 2.5% 2ME (Fig. 1). Furthermore, we examined whether 100 mM DTT can protect RNA form degradation in the lysates of yeast and animal tissue (zebrafish larva). A pellet of liquid-cultured *S. cerevisiae* was homogenized with zirconia beads, and 2 days post fertilization larvae of zebrafish were homogenized with a pestle in Tris-HCl buffers with or without 100 mM DTT, followed by an hour incubation at 22 °C and purification of total RNA. As expected, clear rRNA peaks could be observed in the RNA from the lysates with DTT but not without DTT (Fig. 1). It is noted that the commercial regents for cultured animal cells could not prevent RNA degradation in the lysate of zebrafish (Supplemental Fig.1). We therefore concluded that high concentration of reducing regents can protect RNA form intrinsic RNase.

**Figure 1.**
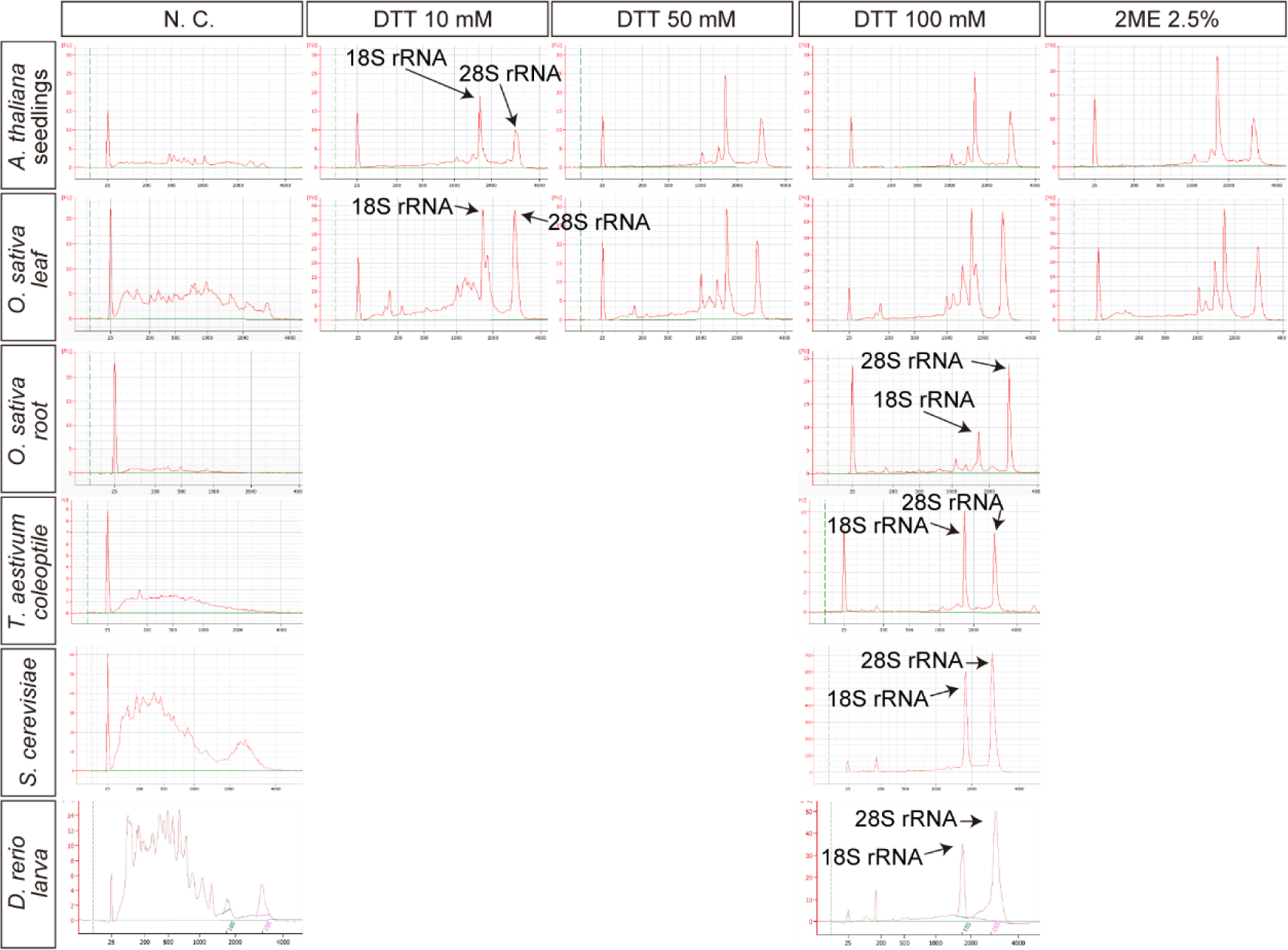
Reducing regents inhibited RNA degradation in plant and animal tissues’ lysate. Bioanalyzer electropherograms of purified RNA from tissue lysates (*A. thaliana* seedlings, *O. sativa* leaves, *O. sativa* root, *T. aestivum* coleoptile, *S. cerevisiae* and *D. rerio*) containing the reducing regents. The columns indicate that the concentrations of the reducing regents in the homogenization buffers. N.C. represents no reducing reagents in buffers (negative control).

### Moderate concentration of reducing regents do not inhibit reverse transcription

Next, we evaluated the efficiency of RT in the presence of the reducing regents. We added several concentrations of DTT or 2ME to RT reaction mixes and performed RT of purified total RNA from *O. sativa*. Then, qPCR for *Os03g0836000* transcript coding for an Actin protein was conducted. Because, in manufacturer’s protocol of RTase, addition of 5 mM DTT is recommended for RT reaction to protect RT enzyme from oxidization, 5 mM DTT is positive control in this experiment. RT products of 5 mM DTT showed 4.4 lower Ct value than the negative control (Fig. 2a). As the concentration of DTT increased, Ct value slightly increased (Fig. 2a). Same tendency was observed for 2ME, but 10 % 2ME drastically diminished the efficiency of RT reaction (Fig. 2a). These results indicate that high concentration of reducing regent (ex. 10% 2ME) inhibits RT reaction but moderate concentration of reducing regent dose not.

**Figure 2.**
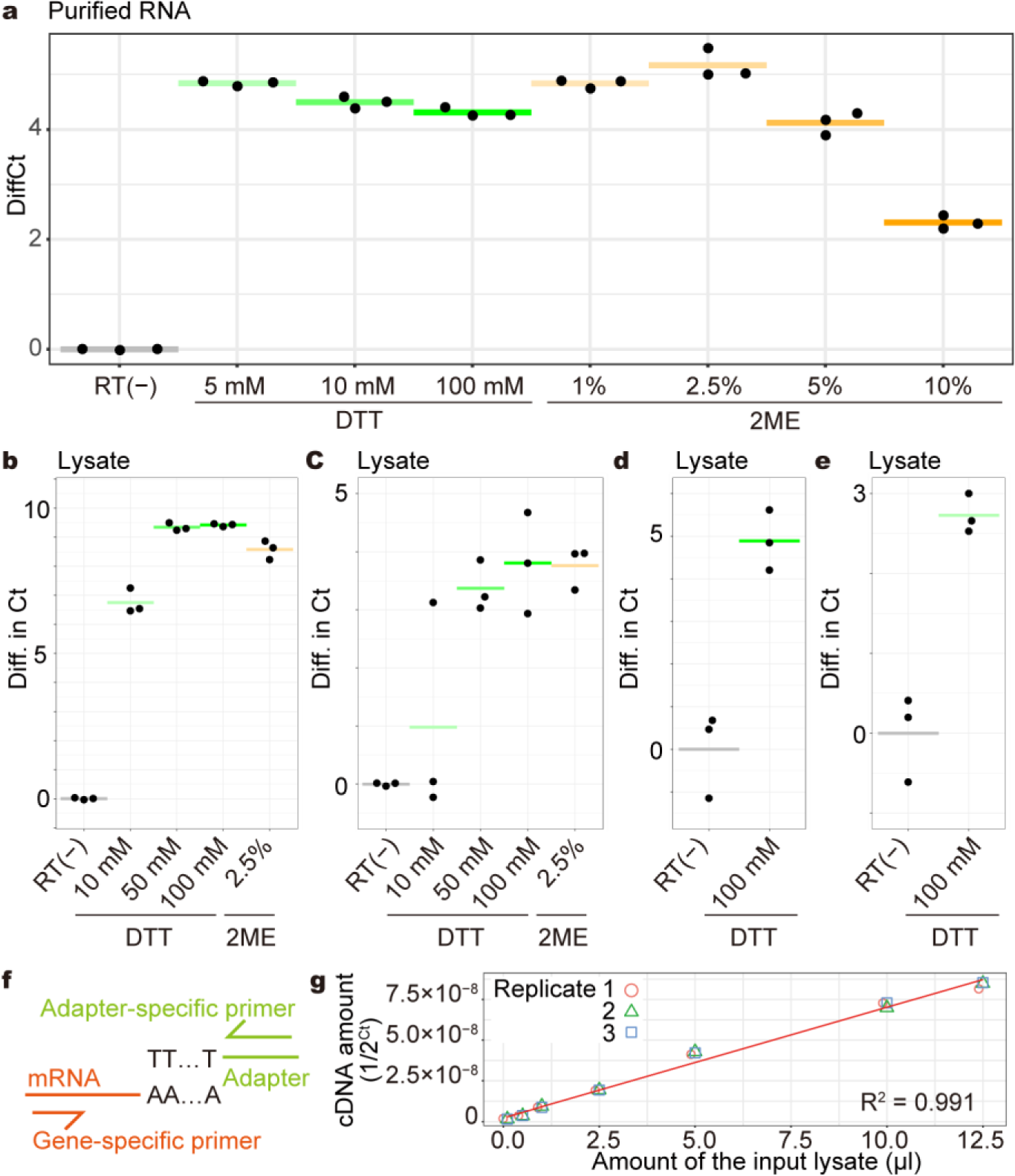
Quantification of cDNA synthesized by direct-lysate reverse transcription. (a) qPCR results for *Os03g0836000* transcript in the cDNA synthesized from mixtures of purified *O. sativa* total RNA and reducing regents. The horizontal axis indicates the concentrations of the reducing regents in the RT reaction mixes. The vertical axis indicates the differences in Ct value compared to the RT(-) negative control. Each point indicates a value of each replicate. Each horizontal line indicates the average of each condition. (b-e) qPCR results for *AT3G18780* transcript (b), *Os03g0836000* transcript (c,d) and *HP620998.1* transcript (e) in cDNA synthesized from lysate of *A. thaliana* seedlings (b), *O. sativa* the youngest fully-expanded leaves (c), *O. sativa* roots (d) and *T. aestivum* coleoptile (e). The horizontal axis indicates the concentrations of the reducing regents in the homogenization buffers. The vertical axis indicates the differences in Ct value compared to the RT(-) negative control. Each point indicates a value of each replicate. Each horizontal line indicates the average of each condition. Same amount of total RNA was used as cDNA template according to RNA concentrations in lysates (250 ng of *A. thaliana* and *O. sativa* or 140 ng of *T. aestivum*). (f) A schematic diagram of the primer sets for cDNA detection. (g) qPCR results for *Os03g0836000* transcript in cDNA synthesized from lysate of *O. sativa* leaves. In the direct-lysate reverse transcription, 0.1, 0.5, 1, 2.5, 5, 10 and 12.5 µL of the lysate were used. Three technical replicates were prepared. The red line is a linear regression line.

To assess whether addition of the reducing regents into homogenization buffer makes it possible to perform RT from plant lysates, we conducted direct-lysate RT of *A. thaliana* seedling and *O. sativa* leaf, root and *T. aestivum* coleoptile, and quantified their actin cDNAs (*AT3G18780*, *Os03g0836000* and *HP620998.1*, respectively) (Fig. 2b-e). Because the genomic DNA remains in the lysates, we conducted RT with dT primers harboring an adapter sequence at the 5’ end (Supplemental Table 1), followed by RT-qPCR with a gene-specific and the adapter-specific primers to detect exclusively cDNA synthesized by the direct-lysate RT (Fig. 2f). Compared to the RT(-) negative controls, amplification of the target cDNAs were observed in the lysates with the reducing regents (Fig. 2b-e). Consistent with RNA degradation levels in the lysates (Fig. 2a), the lysate in the 10 mM DTT buffer showed less cDNA production than the others (Fig. 2b,c). For all plant lysates, the 100 mM DTT buffer showed high production of the actin cDNAs (Fig. 2b-e). In addition, to check the linearity of cDNA synthesis by direct-lysate RT, we conducted direct-lysate RT of varied amounts of the *O. sativa* leaf lysate and quantifies the cDNA with qPCR. The series of direct-lysate RT showed a good linearity between the input amount of the lysate and the cDNA production (R^2^ = 0.991, Fig. 2g). Thus, Direct-RT buffer (100 mM DTT/90 mM Tris-HCl (pH7.6)) was used for the following experiments of direct-lysate RT.

### Direct-lysate RT can produce cDNA comparable to RT from purified RNA

We examined bias in direct-lysate RT comparing with RT of purified total RNA. We compared the results of 3’ RNA-Seq of purified RNA and lysate. The lysate of *O. sativa* leaves was divided into two aliquots. One was purified and used for 3’ RNA-Seq library preparation, whereas the other was directly used for 3’ RNA-Seq library preparation. 29 and 30 technical replicates for cDNA from the purified RNA and lysate were sequenced, respectively. To remove noise derived from total read-number variation, we analyzed 100000 reads subsampled from each technical replicate. The mapping rate onto the transcriptome reference in RNA-Seq of the lysate is approximately 1% higher than that of the purified RNA (Fig. 3a). Also, ratio of rRNA reads of the lysate is approximately 1.7% higher than the purified RNA (Fig. 3b). The numbers of detected genes in the purified RNA and lysate were almost same (Fig. 3c). Comparison of each gene expression level showed that almost all gene were quantified equally in both methods, although 396 of 38194 genes were differentially quantified (Fig. 3d). Among them, 329 genes were quantified as higher amount in the lysate than the purified RNA (Fig. 3d). The transcript length of the highly-quantified genes tended to be short; the average length of the highly-quantified transcripts is 799 nt, while the average length of all transcripts in the reference is 1537 nt (Fig. 3e). There is possibilities that direct-lysate RT could efficiently produce cDNA of short transcripts in the lysate of *A. thaliana* seedlings and/or that some of short transcripts might be diminished through RNA purification process. Also, we conducted same comparison of RNA-Seq of lysate and purified RNA from budding yeast and zebrafish larva. In the case of the yeast, among 6713 genes, 37 and 48 genes were highly and lowly quantified in RNA-Seq of the lysate (Supplemental Fig. 2a). On the other hand, in the case of the zebrafish larva, among 57775 genes, 6913 and 1048 genes were highly and lowly quantified in RNA-Seq of the lysate (Supplemental Fig. 2b). Unlike *A. thaliana* lysate, any bias in the transcript length was not observed in both yeast and zebrafish lysates (Supplemental fig. 2c and d). We concluded that direct-lysate RT method can produce comparable, or even higher, amount of cDNA to that synthesized from purified RNA, although RT efficiencies of some transcripts could be specifically altered by the composition of lysates.

**Figure 3.**
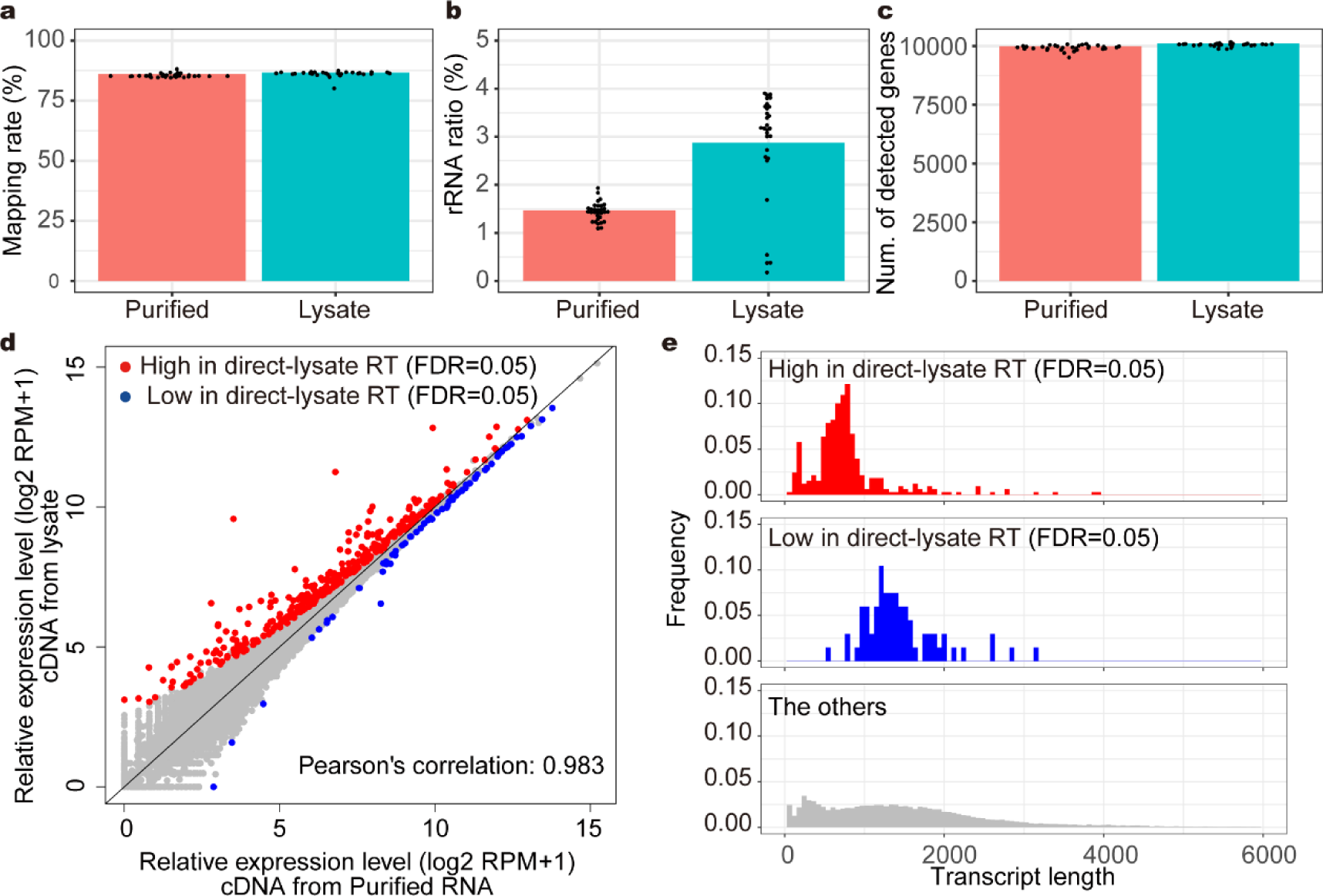
Direct-lysate reverse transcription could generate cDNA comparable to reverse transcription of purified RNA. The results of 3’ RNA-Seq of purified RNA and lysate of *A. thaliana* seedlings. 100000 subsampled reads for each technical replicate were used for the following analysis. (a) Boxplot of mapped read ratio to 100000 reads. (b) Boxplot of the rRNA read ratio to 100000 reads. (c) Boxplot of the number of detected genes. Each point indicates value of each technical replicate. (d) Scatter plot of log2 RPM+1 of each gene. Average of technical replicates was plotted. Differentially quantified genes (DQGs) were detected between RNA-Seq for purified RNA and lysate (FDR = 0.05). (e) Transcript length distribution of DQGs with higher rpm values in the method using lysate and the other genes. The histograms from 0 nt to 6000 nt are shown.

### Direct-lysate targeted RNA-Seq (DeLTa-Seq)

To develop a cost-effective and accurate gene quantification method applicable for using direct-lysate RT, we tried to take advantage of targeted amplicon RNA-Seq^29^. We designed the gene-specific primers for all *A. thaliana* genes (googlesite URL). Here, we focused on the 100 marker genes focused on in the previous study^33^. We integrated the PCR-based targeted RNA-Seq with direct-lysate RT, named “**D**ir**e**ct-**L**ysate **Ta**rgeted RNA-Seq (DeLTa-Seq) (Fig. 4a); First, 1^st^ index was added by the direct-lysate RT with oligo-dT primer harboring 1^st^ indexes, following by sample pooling (up to 384 samples). The RT primer contains unique molecular identifier (UMI). Second, a part of the adapter sequence was added to the cDNA of target genes using gene-specific primers harboring a part of the adapter sequence. Third, 2^nd^ indexing and target amplification were conducted with PCR. If needed, up to 384 libraries can be pooled, which enables us to sequence up to 147456 samples at once. To evaluate target enrichment by DeLTa-Seq, we prepared two libraries, non-targeted RNA-seq and targeted RNA-Seq (DeLTa-Seq) from the same lysate of *A. thaliana* seedling, followed by sequencing with MiSeq, resulting in 6694751 reads of the non-targeted RNA-Seq and 203450 reads of DeLTa-Seq. All target genes are quantified more highly in targeted RNA-Seq than in non-targeted RNA-Seq (Fig. 4b). The ratio of reads corresponding to the target genes increased from 0.45% in the non-targeted RNA-Seq to 94.9% in the targeted RNA-Seq; more than 200-fold-enrichment for the target cDNA was achieved even for direct-lysate RT (Fig. 4c).

**Figure 4.**
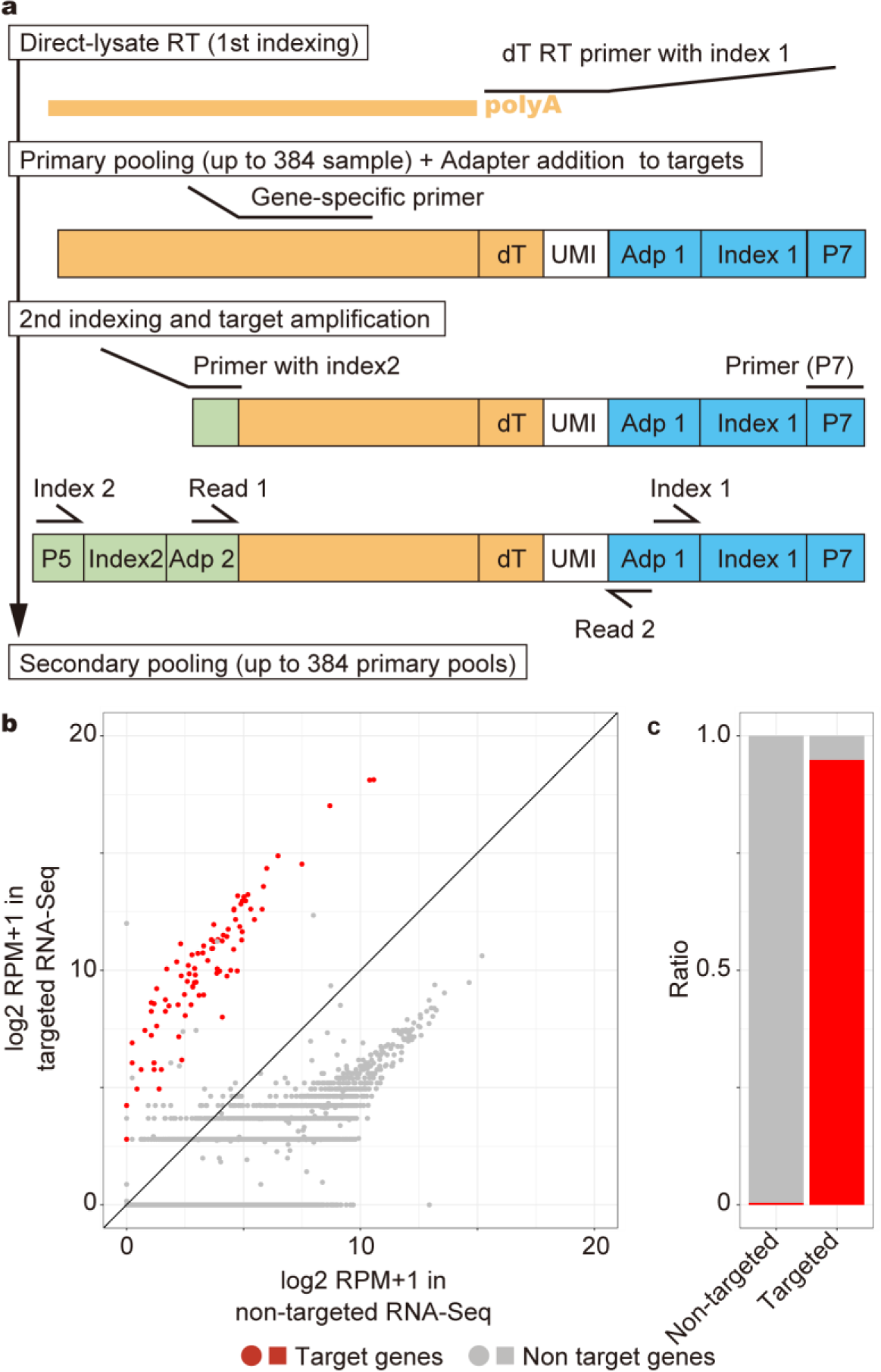
DeLTa-Seq works for *A. thaliana* seedlings. (a) A schematic diagram of DeLTa-Seq library preparation. P7 and P5 are adapter sequences binding with the oligonucleotides on the flow cells supplied by illumina Adp 1 and Adp 2 are adapter sequences for the sequencing. UMI indicates unique molecular identifier. (b) Scatter plot of log2 RPM+1 values of each gene in targeted RNA-Seq and non-targeted RNA-Seq of *A. thaliana* seedlings’ lysate. (c) The bar graph represents the ratio of reads correspond to target and non-target genes in targeted RNA-Seq and non- targeted RNA-Seq.

### Targeted RNA-Seq works with low amount of input RNA

Next, we examined the required amount of input RNA for the targeted RNA-Seq. We prepared targeted RNA-Seq libraries from 0.01 ng, 0.1 ng, 1 ng and 10 ng of a same purified *A. thaliana* total RNA with 96, 96, 10 and 5 technical replicates, respectively. After sequencing with MiSeq, 10,000 reads were subsampled from each replicate. The mean ratios of the target reads to the subsampled reads were 89.5%, 87.1% and 87.3% in the libraries of 0.1, 1 ng and 10 ng inputs, respectively, whereas 1.31% in the libraries of 0.01 ng input (Fig. 5a). While 86 genes were totally detected, the number of detected target genes was saturated in the libraries of more than 0.1 ng input (69.6 and 74.4 genes on average in 1 ng and 10 ng), whereas very few target genes (1.26 genes on avarage) were detected in the libraries of 0.01 ng input (Fig. 5b). Pearson’s correlation coefficient of log2 RPM+1 of the target genes in each library to each 10-ng-input library was also saturated in the libraries of more than 0.1 ng input (Fig. 5c). Based on these results, targeted RNA-Seq roughly requires 1 ng input of total RNA, and would be applicable for small tissues.

**Figure 5.**
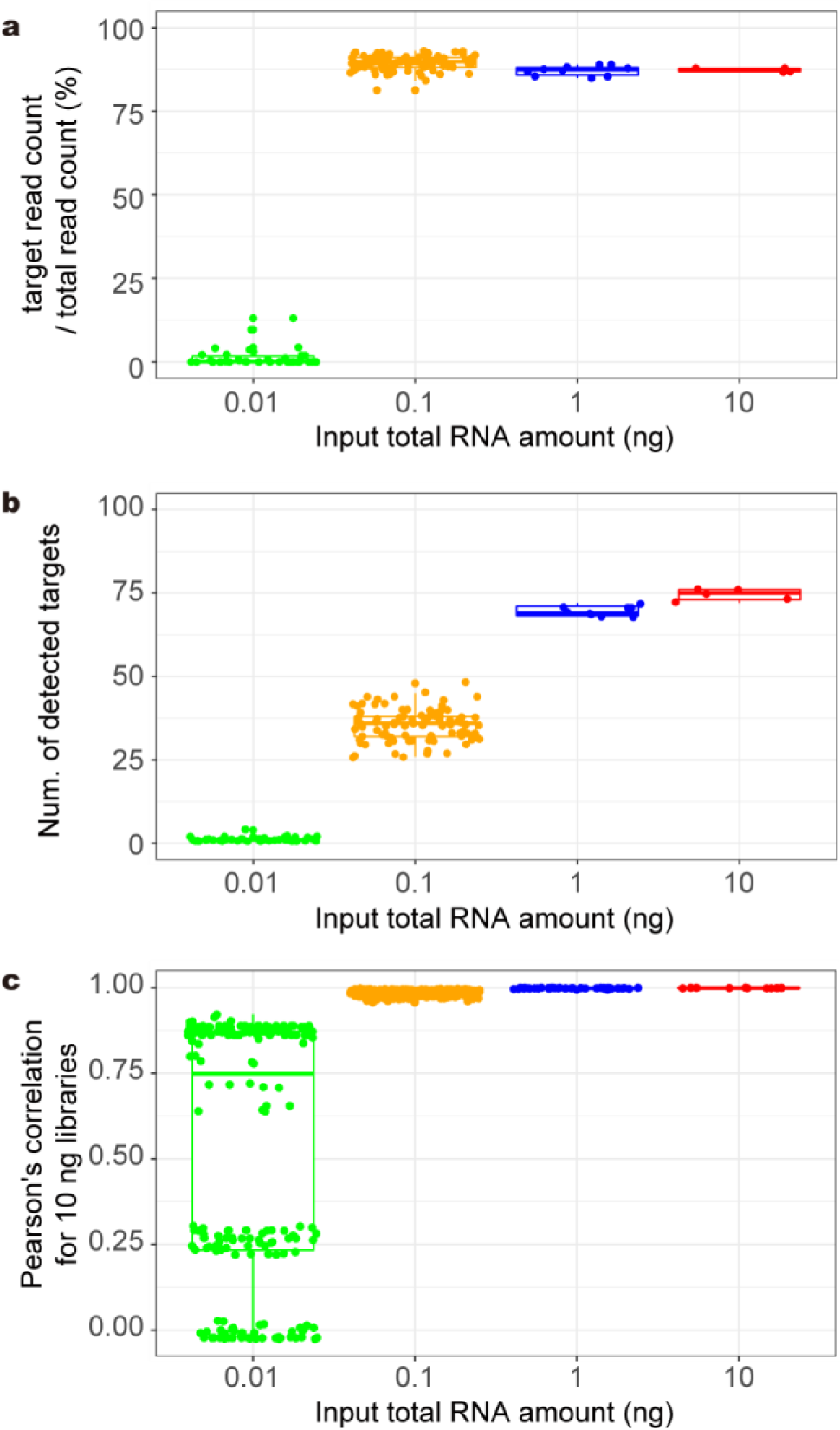
Required amount of input RNA for the targeted RNA-Seq. (a-c) The result of targeted RNA-Seq using 0.01 ng (n = 96), 0.1 ng (n = 96), 1 ng (n = 10) and 10 ng (n = 5) of purified *A. thaliana* RNA. 10000 reads subsampled from each library was used for the analyses. Boxplots of the ratio of target reads per total read (a), the numbers of detected target genes (b) and Pearson’s correlation coefficients to each results of 10ng RNA. (c). Each point indicates value of each technical replicate.

### Targeted RNA-Seq improved reproducibility and cost of gene quantification

To examine the reproducibility and performance of mRNA quantification by targeted RNA-Seq, we prepared 96 technical replicates of targeted and non-targeted RNA-Seq libraries, respectively, using single tube of RNA of *A. thaliana* seedlings and sequenced them with HiSeq. In our targeted RNA-Seq, we used 10-bases unique molecular identifiers (UMI), which can theoretically distinguish a million of RNA molecules of each target transcript^31, 34^. All reads were used after UMI preprocessing, resulting in 96 technical replicates of non-targeted RNA-Seq of 1786180 UMI counts on average and of targeted RNA-Seq of 344215 UMI counts on average. UMI conversion efficiencies were 0.805 on average in non-targeted RNA-Seq, and 0.434 on average in targeted RNA-Seq, respectively (Supplemental fig. 3). We merged all UMI counts of targeted and non-targeted RNA-Seq (33044648 and 171473284 UMI counts, respectively) to be used as ‘full data’. First, to evaluate reproducibility of our targeted RNA-Seq protocol, we subsampled from 10000 to 150000 UMI counts from each technical replicate of targeted RNA-Seq. Then, Pearson’s correlation coefficients of log2 RPM+1 of the target genes in each subsampled data set for the full data were calculated. With 150000 UMI counts, 96 replicates showed high reproducibility (the average of Pearson’s correlation coefficients for the full data: 0.977 ± 0.0039 [mean ± S.D.]) (Fig. 6a). Next, to examine quantification performance, we merged every eight technical replicates of non-targeted and targeted RNA-Seq into one, resulting in 12 pseudo-technical replicates, respectively. Then, we subsampled from 1000 to 2000000 UMI counts of targeted RNA-Seq and from 10000 to 10000000 UMI counts of non-targeted RNA-Seq from each pseudo-technical replicate. Reproducibility of quantification of each target gene was almost saturated at 250000 UMI counts in the targeted RNA-Seq, while 5000000 UMI counts in the non-targeted RNA-Seq (Fig. 6b). Similarly, Pearson’s correlation coefficients of log2 RPM+1 of the target genes in each subsampled data set for the full data was saturated at 250000 and 5000000 UMI counts in the targeted and non-targeted RNA-Seq, respectively (Fig. 6c). Thus, targeted RNA-Seq (DeLTa-Seq) would enable us to conduct cost-effective and accurate quantification of target gene expression by reducing sequencing cost.

**Figure 6.**
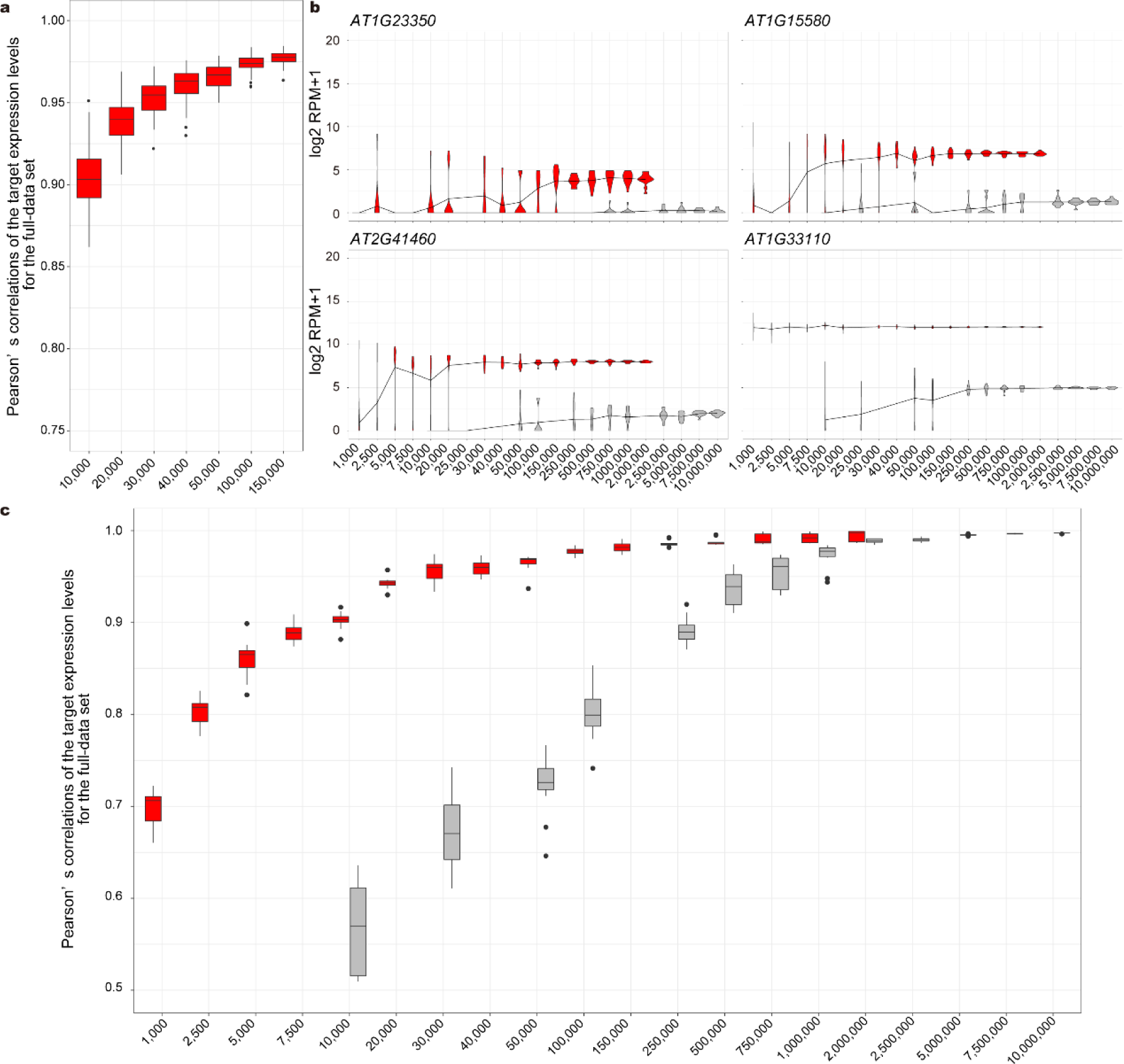
Reproducibility and required UMI counts for the targeted and non-targeted RNA-Seq. (a) A plot of Pearson’s correlation coefficients of the target log2 RPM+1 between each subsampled 96 technical replicates and the full-dataset of targeted RNA-Seq. The replicates were prepared using single tube of purified RNA of *A. thaliana* seedlings. (b) Plots of examples of the target log2 RPM+1 in each size of subsampled sets from 12 pseudo-technical replicates of targeted and non-targeted RNA-Seq results. The same purified RNA was used for the preparation of 96 technical replicates of non-targeted RNA-Seq. Then, eight RNA-Seq results were merged, resulting in 12 pseudo-technical replicates. (c) Plot of Pearson’s correlation coefficients of the target log2 RPM+1 between each subsampled set from the 12 pseudo-technical replicates and the full-dataset of targeted and non-targeted RNA-Seq, respectively. (a-c) Red indicates targeted RNA-Seq. Gray indicates non-targeted RNA-Seq. horizontal axes indicate the numbers of subsampled UMI counts.

## Discussion

Direct-lysate RT are utilized in analyses of cultured animal cells to reduce cost and labor for RNA purification, followed by gene expression analysis such as RT-qPCR and RNA- Seq^21, 35^. In these direct-lysate RT methods, RNase inhibitors are usually added to the lysis buffers to inhibit RNA degradation by intrinsic RNase. Because commercial RNase inhibitors are expensive, it is not realistic to use them in preparation of lysates from large size and/or large number of crude plant and animal tissues. Notably, we demonstrated that these methods cannot be applicable to plant and animal tissues (Supplemental fig. 1). In this study, we developed Direct-RT buffer that realizes cost-effective direct-lysate RT of crude plant, yeast and animal samples. DTT could reduce disulfide bonds required for the activity of RNase in lysates^24^. Because no loss of RNA in purification occurs through direct-lysate RT, this technique is beneficial also for gene expression analysis in small tissues from which it is difficult to purify enough amount of RNA for the experiments. Furthermore, DTT may contribute long-term maintenance of endogenous RNase inhibitor activity under storage at -20 °C^23, 36^. Thus, our Direct-lysate RT technique will be useful not only for gene expression analysis but also RNA storage.

Comparison of the non-targeted RNA-Seq results of the purified RNA and the lysate revealed that some transcripts were differentially reverse transcribed in the direct-lysate RT than in RT of the purified RNA (Fig. 3d and Supplemental fig. 2). For example, in the case of *A. thaliana* seedlings, the amount of cDNA of rRNA was twice in the direct-lysate RT as much as in RT with the purified RNA, but less than 4% (Fig. 3b). Previous studies showed that unspecific priming from A-rich region of rRNA with dT primer occurs in RT step^19, 37^. Considering that lysate will contain higher concentration of salt than purified RNA solution and that salt increases the stability of base-paring of nucleic acid^38^, unspecific priming by dT primer against non-polyA RNA such as rRNA would occur in direct-lysate RT more than in RT with purified RNA. Also, direct-lysate RT in the lysate of *A. thaliana* seedlings might produce more cDNA of relatively short transcripts (Fig. 3e). Given that numerous transcripts were differentially quantified in the lysate of zebrafish (Fig. S2), there is another possibility that the composition of the lysates affected the binding between transcripts and the magnetic beads, causing biases in RNA purification step. It is recommended to compare the RNA-Seq of a lysate with the RNA-Seq of purified RNA as a pilot experiment.

In this study, we examined how much input are required for accurate gene quantification. At least, 0.1 ng total RNA per sample was required for stable quantification of the target genes (Fig. 5b,c), but targeted-RNA-Seq with 0.1 ng total RNA resulted in abundant useless reads (Fig. 5a). The amount of the 100 target RNAs in 0.01 ng total RNA could be less than 0.1 pg (1% of total RNA) equal to mRNA amount of one cell^39^, resulting in few detected targets (Fig. 5b) as observed in single cell RNA-Seq. Dropouts is known as the phenomenon in which single-cell RNA-Seq can capture only small fraction of the transcriptome of each cell and the count of most genes are zero^40^. Considering that one cell contains 10 pg of total RNA^39^, the requirement of DeLTa-Seq will be acceptable for small tissue sample consisting of roughly 100 cells.

A high resolution of time, space, perturbation, and stochastic variation should be required to understand complex biological systems via transcriptome analysis. From the viewpoint of sequencing cost, it is difficult to apply non-targeted RNA-Seq strategy to thousands of samples. Although the required read counts for robust quantification by targeted RNA-Seq depends on expression levels of target transcripts, our study indicated that 150000 UMI counts could be enough for targeted RNA-Seq to quantify target transcripts (Pearson’s correlation coefficient > 0.98) (Fig. 6). Given the high throughput of recent illumina DNA sequencers and our UMI conversion efficiencies (Supplemental Fig. 3), we can sequence thousands of samples per lane, resulting in cost-effective quantification of genes of interest. For example, with HiSeq 4000, which can generate 300 million reads per lane, two thousand samples can be sequenced at a time. Coupled with skipping RNA purification step, DeLTa-Seq can drastically reduce cost and labor in mRNA quantification. DeLTa-Seq is useful not only for large scale studies handling thousands of samples such as field transcriptome and chemical screening but also in-depth studies of specific biological systems. For example, a researcher will conduct limited number of comprehensive RNA-Seq, and find genes of interest. DeLTa-Seq can measure expressions of the genes of interest in many samples required for in-depth analysis. Notably, in our protocol, same cDNA can be shared for both non-targeted and targeted RNA-Seq. This advantage enables us to select non-targeted and targeted RNA-Seq according to the aim of each round. DeLTa-Seq will realize cost-effective and accurate quantification of these genes in numerous samples. DeLTa-Seq can be applicable for animals as well as plants. This technique will open up a novel field of both plant and animal biology. (3522/5000 words)

## Materials and Methods

### Plant materials

Seeds of *Oryza sativa* L. *japonica* ‘Koshihikari’ were sown in nursery trays followed by cultivation for two weeks under 16-h light/8-h dark and 30°C/ 20°C cycles.

Seeds of *Arabidopsis thaliana* L. (Col-0, CS70000) were sterilized in 90% EtOH. Then, they were sown on 1/2× Murashige and Skoog medium with 0.25% gellan gum (Wako, Osaka, Japan), 0.05% (v/v) PPM-100 (Plant Cell Technology, Washington, D.C., USA), 1.25 mM MES-KOH (pH 5.7) and 0.5% sucrose following by cultivation for 21 days at 20°C under 16-h light/8-h dark cycles.

Seeds of *Triticum aestivum* L. ‘Chinese Spring’ were sown on 200 μL zirconia beads YTZ-1 (AS-ONE, Osaka, Japan) with 110 μL nuclease-free water in 2 mL microtubes LT-0200 (INA OPTICA, Tokyo, Japan) followed by cultivation for 3 days under 24-h dark and 4°C and 8 days under 24-h light and 20°C.

### Budding yeast material

*S. cerevisiae* BY4742 strain was cultured in Synthetic Complete (SC) medium at 28 °C. The yeast in the exponential phase was used in this study.

### Animal material

Fertilized eggs of zebrafish (*Danio rerio*) were obtained by cross of lab stock AB provided by Zebrafish International Resource Center (ZIRC) located at the University of Oregon. They were maintained at 28 °C under dark condition till sampling. Larvae were sacrificed at 48 hours post fertilization.

### Preparation of purified *O. sativa* and *A. thaliana* RNA

At two weeks after seeding, the youngest fully expanded leaves of *O. sativa* were frozen in liquid nitrogen, and stored at -80 °C until RNA isolation for RNA-Seq. Frozen samples were homogenized with zirconia beads YTZ-4 (AS-ONE, Osaka, Japan) using TissueLyser II (Qiagen, Hilden, Germany), and total RNA was then extracted using the Maxwell 16 LEV Plant RNA Kit (Promega, Madison, WI, USA) and Maxwell 16 Automated Purification System (Promega). The concentration of RNA was measured using QuantiFluor RNA System (Promega) and Quantus Fluorometer (Promega).

Seven days after seeding, bulked seedlings of *A. thaliana* were homogenised with zirconia beads YTZ-4 and TissueLyser II (Qiagen, Hilden, Germany), and total RNA was then extracted using the Maxwell 16 LEV Plant RNA Kit (Promega, Madison, WI, USA) and Maxwell 16 Automated Purification System (Promega, Madison, WI, USA).

The concentration of RNA was measured using Quantus Fluorometer (Promega, Madison, WI, USA).

### Preparation of lysate

Seedlings of *A. thaliana* (Max 9 seedlings) were homogenised with zirconia beads YTZ- 4 and TissueLyser II in 400 µl of 90 mM Tris-HCl (pH 7.6) containing reducing regents (10, 50 or 100 mM DTT or 2.5% 2ME), CL buffer [50 mM Tris-HCl pH 7.6, 0.1% Igepal CA-630, 150 mM NaCl and 1 mM DTT]^23^ or CellAmp Processing Buffer (TaKaRa, Kusatsu, Japan) or Lysis Solution of SuperPrep (TOYOBO, Osaka, Japan). DTT solution should be freshly prepared or stored at -20 °C. According to the manufacture’s manual, for SuperPrep, 76 µl Stop solution (TOYOBO) and 4 µl RNase Inhibitor (TOYOBO) were added after incubation for 5 min at 22 °C.

Each youngest fully expanded leaf of *O. sativa* was frozen in liquid nitrogen, and stored at -80 °C until homogenization. Each frozen sample was homogenized with zirconia beads YTZ-4 using TissueLyser II. Then, 400 µl of the same buffers used for *A. thaliana* seedlings were added, and, immediately, the samples were mixed well by vortexing. According to the manufacture’s manual, for SuperPrep, 76 µl Stop solution and 4 µl RNase Inhibitor were added 5 min after incubation at 22 °C.

Each coleoptile with first leaf of *T. aestivum* was frozen in liquid nitrogen, and stored at -80 °C until homogenization. Each frozen sample was homogenized with the zirconia beads YTZ-4 using TissueLyser II. Then, 400 µl of the Direct-RT buffer was added, and, immediately, the samples were mixed well by vortexing.

*S. cerevisiae* in the exponential phase was homogenised with zirconia beads YTZ-1 and MS-100R (TOMY, Tokyo, Japan) in 350 µl of 90 mM Tris-HCl (pH 7.6) containing 100 mM DTT.

Three *D. rerio* larvae were homogenized in 50 µl of the same buffers used for *A. thaliana* seedlings with Biomasher Ⅱ (Nippi, Tokyo, Japan). According to the manufacture’s manual, for SuperPrep, 9.5 µl Stop solution and 0.5 µl RNase Inhibitor were added after incubation for 5 min at 22 °C.

The concentration of RNA in the lysates were measured using the Quant-iT RNA Assay Kit (Thermo Fisher Scientific, Waltham, MA, USA) and Infinite M1000 PRO (TECAN, Zurich, Switzerland).

### Purification of RNA from lysates

Lysate was incubated for an hour at 22 °C. Then, 2.5× volume of AMpure XP beads (Beckman Coulter, Brea, CA, USA) was added to a lysate. The mixture was purified following the manufacturer’s instructions. The RNA was then eluted with the equal volume of RNase-free water to the lysate. One microliter of the purified RNA solution was used for electrophoresis using a Bioanalyzer 2100 with Agilent RNA nano or pico kit (Agilent Technologies, Santa Clara, CA, USA) to check for the quality.

### Direct-lysate reverse transcription

RT reactions were conducted with 1 μL of 2 μM RT primer (CAGAAGACGGCATACGAGATGCGTCTACGTGACTGGAGTTCAGACGTGTGC TCTTCCGATCNNNNNNTTTTTTTTTTTTTTTTTTV), 0.4 μL of 25 mM Dntp (Advantage UltraPure dNTP Combination Kit) (TaKaRa), 4.0 μL of 5× SSIV Buffer (Thermo Fisher Scientific), 0.1 μL of SuperScript IV reverse transcriptase (200 U/μL, Thermo Fisher Scientific), lysates and nuclease-free water to make a volume of 20 μL. Reverse transcription was carried out at 65°C for 10 min, following by incubation at 80°C for 15 min to inactivate the enzyme.

### Quantitative PCR analysis of cDNA (RT-qPCR) for figure 2

250 ng (*A. thaliana* and *O. sativa*) or 140 ng (*T. aestivum*) of total RNA in lysate was used for RT. Then, the reaction mix of *A. thaliana* and *O. sativa* was diluted with nucleases-free water by 20 times. The reaction mix of *T. aestivum* was used for the following experiments without dilution. The cDNA amount of *AT3G18780*, *Os03g0836000* or *HP620998.1* was measured by qPCR analysis. The composition of the qPCR mixture is described below; diluted cDNA solutions 2 µl, gene-specific primer (CTTGCACCAAGCAGCATGAA for *AT3G18780* or GTGTGTCGGTACTTTCGTCG for *Os03g0836000*, TGACCGTATGAGCAAGGAG for *HP620998.1*^41^, P7 primer for the adapter sequence added in the RT step (CAAGCAGAAGACGGCATACGAGAT) and 5 µl of KAPA SYBR FAST qPCR Master Mix (2×) (KAPA BIOSYSTEMS, Wilmington, MA, USA). qPCR was conducted using LightCycler 480 System II (Roche Diagnostics, Basel, Switzerland) with the following program: enzyme activation at 95 °C for 3 min, 40 cycles at 95 °C for 3 s and 60 °C for 30 s for amplification.

### Non-targeted RNA-Seq for figure 3

Non targeted RNA-Seq were conducted according to Lasy-Seq ver. 1.1 protocol^19^ (https://sites.google.com/view/lasy-seq/). RT reactions were conducted with 1 μL of 2 μM of a RT primer (CAGAAGACGGCATACGAGATxxxxxxxxGTGACTGGAGTTCAGACGTGTGCTC TTCCGATCNNNNNNTTTTTTTTTTTTTTTTTTVT, xxxxxxxx indicate index sequences for multiplex sequencing. See supplemental table 1), 0.4 μL of 25 mM dNTP, 4.0 μL of 5× SSIV Buffer, 2.0 μL of 100 mM DTT, 0.1 μL of SuperScript IV reverse transcriptase, template RNA (lysate or purified RNA solution) and nuclease-free water to make a volume of 20 μL. Reverse transcription was carried out at 65°C for 10 min, following by incubation at 80°C for 15 min to inactivate the enzyme. Then, all RT mixture of samples were pooled and purified with Wizard SV Gel and PCR Clean-Up System (Promega, Madison, WI, USA) according to the manufacture’s manual except for using 80% EtOH instead of Membrane Wash Solution. The purified cDNA was eluted with 30 μL of nuclease-free water. Second strand synthesis was conducted on the pooled samples (30 μL) with 4 μL of 10× blue buffer (Enzymatics, Beverly, MA, USA), 2 μL of 2.5 mM dNTP (Advantage UltraPure dNTP Combination Kit), 1 μL of 100 mM DTT, 1 μL of RNaseH (5 U/μL, Enzymatics), 2 μL of DNA polymerase I (10 U/μL, Enzymatics). The reaction was conducted at 16°C for 2 h and kept at 4°C until the next reaction. To avoid the carryover of large amount of rRNAs, RNase treatment was conducted on the mixture with 1 µL of RNase T1 (1 U/µL, Thermo Fisher Scientific) and 1 µL of RNase A (10 ng/µL, NIPPON GENE, Tokyo, Japan). The reaction was conducted at 37°C for 5 min and 4°C until the next reaction. Then, purification was conducted with 0.8× volume of AMpure XP beads according to the manufacturer’s manual, following by elution with 36 µL nuclease-free water. Fragmentation, end-repair and A-tailing were conducted in the mixture of 17.5 µL of the dsDNA, 2.5 µL of 10× Fragmentation Buffer (Enzymatics, Beverly, MA, USA) and 5 µL of 5× WGS Fragmentation Mix (Enzymatics, Beverly, MA, USA). The mixture was kept at 4°C before the following reaction. The reaction was conducted at 4°C for 1 min, 32°C for 7 min, 65°C for 30 min and 4°C until the next reaction. Adapter for the next ligation step was prepared by annealing 100 mM of A*C*C*GAGATCTACACACTCTTTCCCTACACGACGCTCTTCCGA*T*C*T and /5Phos/G*A*T*CGGAAGAGCGTCGTGTTAAATGTA*T*A*T (* signifies a phosphorothioate bond. /5Phos/ signifies a phosphorylation.) using a thermal cycler with the following program: 95°C for 2 min, slow-cooled to 45°C (0.1°C/s), followed by 45°C for 5 min. 10 µL of 5× Ligation buffer (Enzymatics), 8 µL of nuclease-free water, 2 µL of 4 mM annealed adapter were added to the above reaction mix, following by adding 5 µL of T4 DNA ligase (Enzymatics). The ligation was conducted at 20°C for 15 min. The adapter-ligated DNA was purified with 0.8× volume of AMpure XP beads, twice, followed by elution with 17 µL of nuclease-free water. To optimise PCR cycle for library amplification, qPCR was conducted with 3.5 µL of the adapter-ligated DNA, 5 µL of KAPA HiFi HotStart ReadyMix (KAPA BIOSYSTEMS), 0.5 µL of Evagreen, 20× in water (Biotium, Fremont, CA, USA), 0.5 µL of 10 µM 5× WGS_Pr_R1-5’ primer (AATGATACGGCGACCACCGAGATCTACACTCGTCGGCAGCGTC) and 0.5 µL of 10 µM PCR_P7 primer (CAAGCAGAAGACGGCATACGAGAT). Reaction was carried out using LightCycler 480 II at 95°C for 5 min, 30 cycles of 98°C for 20 s, 60°C for 15 s, 72°C for 40 s, followed by 72°C for 3 min, then held at 4°C. The optimal number of the PCR cycle was the cycle corresponding to the middle of exponential phase. Then, library was amplified with 12 µL of the adapter-ligated DNA, 15 µL of KAPA HiFi HotStart ReadyMix, 1.5 µL of 10 µM 5× WGS_Pr_R1-5’ primer and 1.5 µL of 10 µM PCR_P7 primer. Reaction was carried out at 95°C for 5 min, the optimized cycles of 98°C for 20 s, 60°C for 15 s, 72°C for 40 s, followed by 72°C for 3 min, then held at 4°C. The amplified library was purified with equal volume of AMpure XP beads and eluted with 15 µL of nuclease-free water. One microliter of the library was used for electrophoresis using a Bioanalyzer 2100 with Agilent High Sensitivity DNA kit to check for the quality. Sequencing of 50-bp single-ends using HiSeq (Illumina, San Diego, CA, USA) was carried out.

### The design of gene-specific primers for *A. thaliana*

We designed gene-specific primers for *A. thaliana* cDNA sequences using primer3^42^. The parameters were as below; Concentration of Monovalent Cations : 40 mM, Concentration of Divalent Cations: 6mM, Annealing Oligo Concentration: 25nM, Primer Size Min/Opt/Max : 20/23/25, Primer Tm Min/Opt/Max : 60/65.5/70, Primer GC% Min/Opt/Max : 30/50/55, Max Tm Difference : 5, Product Size Ranges : 289-889, Number To Return : 5,PRIMER_RIGHT_INPUT : CAAGCAGAAGACGGCATACGAGAT, PRIMER_MAX_END_STABILITY. Then, candidates of gene-specific primers were concatenated as below; TCGTCGGCAGCGTCAGATGTGTATAAGAGACAG+gene-specif-perimer. All candidates of gene-specific primers for all *A. thaliana* genes are available in google site URL.

### Direct-lysate targeted RNA-Seq (DeLTa-Seq) for figure 4

RT reactions and sample-pooling were conducted according to the above protocol for the non-targeted RNA-Seq, following by elution of cDNA with 40 µl of nuclease-free water. 20 μL of the pooled cDNA was used. RNase treatment was conducted on the mixture with 1 µL of RNase T1 and 1 µL of RNase A. The reaction was conducted at 37°C for 5 min and 4°C until the next reaction. Then, purification was conducted with 0.8× volume of AMpure XP beads according to the manufacturer’s manual, following by elution with 10 µL nuclease-free water. The adapter sequence was added to the target transcripts in the library amplification step. The reaction was conducted as below; 4 µL of the RNase-treated cDNA solution, 5 µL of KAPA HiFi HotStart ReadyMix and 1 µL of gene-specific primer mix (total 100 µM, See Supplemental Table 2). The reaction was conducted at 95°C for 5 min, 98°C for 20 sec, 60°C for 15 sec, 72°C for 40 sec 5 min and 4°C until the next reaction. Then, purification was conducted with 0.8× volume of AMpure XP beads according to the manufacturer’s manual, following by elution with 12 µL nuclease-free water. To optimise PCR cycle for library amplification, qPCR was conducted with 3.5 µL of the adapter-added DNA, 5 µL of KAPA HiFi HotStart ReadyMix, 0.5 µl of Evagreen, 20× in water, 2^nd^-index primer (AATGATACGGCGACCACCGAGATCTACACxxxxxxxxACACTCTTTCCCTACAC GA, xxxxxxxx indicates index sequences for multiplex sequencing. See supplemental table 3) and 0.5 µL of 10 µM PCR_P7 primer (CAAGCAGAAGACGGCATACGAGAT). Reaction was carried out using LightCycler 480 II at 95°C for 5 min, 30 cycles of 98°C for 20 s, 60°C for 15 s, 72°C for 40 s, followed by 72°C for 3 min, then held at 4°C. The optimal number of the PCR cycle was the cycle corresponding to the middle of exponential phase. Then, library was amplified with 7 µL of the adapter-added DNA, 10 µl of KAPA HiFi HotStart ReadyMix, 1 µL of 10 µM 2^nd^ index primer (Supplemental Table 2) and 1 µL of 10 µM PCR_P7 primer. Reaction was carried out at 95°C for 5 min, the optimized cycles of 98°C for 20 s, 60°C for 15 s, 72°C for 40 s, followed by 72°C for 3 min, then held at 4°C. The amplified library was purified with equal volume of AMpure XP beads and eluted with 15 µL of 10 mM Tris-HCl (pH 7.6). One microliter of the library was used for electrophoresis using a Bioanalyzer 2100 with Agilent High Sensitivity DNA kit to check for the quality. Sequencing of 140-bp single-ends was carried out with MiSeq (Illumina, San Diego, CA, USA). The sequences of the first 50-bp of 5’-end were used in the following analysis.

An update of DeLTa-Seq protocol is available in https://xxxxxx.

### Targeted RNA-Seq for figure 5

RNA was extracted from bulked homogenate of approximate 10 seedlings of *A. thaliana* in 14 days after sowing. Then, 0.01 ng, 0.1 ng, 1 ng and 10 ng of the purified RNA were used for the reverse transcription. The number of technical replicates were 96, 96, 10 and 5, respectively. The following experiments were conducted as described in the section of ‘Direct-lysate targeted RNA-Seq (DeLTa-Seq) for figure 4’.

### Preparation and sequencing of 96 technical replicates of non-targeted and targeted RNA-Seq for Figure 6

RNA was extracted from bulked homogenate of approximate 250 seedlings of *A. thaliana* in 7 days after sowing. RT reactions, pooling and purification were conducted as described in the section of ‘Non-targeted RNA-Seq for figure 3’ except for the amount of template RNA (250 ng) for each technical replicate. The pooled cDNA was eluted with 50 μL of nuclease-free water in the purification step.

For non-targeted RNA-Seq, 20 μL of the pooled cDNA was used. The following steps were conducted as described in the section of ‘Non-targeted RNA-Seq for figure 3’ with some changes; dsDNA after RNase treatment was eluted with 10 µL nuclease-free water. The optimization of PCR amplification cycle were conducted with 4 µL of the adapter-ligated dsDNA. Then, library was amplified using 7 µL of the adapter-ligated DNA. The amplified library was purified and eluted with 15 µL of 10 mM Tris-HCl (pH 7.6).

For targeted RNA-Seq, the following experiments were conducted as described in the section of ‘Direct-lysate targeted RNA-Seq (DeLTa-Seq) for figure 4’.

Equal amount by mole of non-targeted and targeted RNA-Seq library were mixed based on qPCR-based quantification with Kapa Library Quantification Kit (KAPA BIOSYSTEMS, Wilmington, MA, USA). Sequencing of 150-bp paired-ends using HiSeq was carried out.

### Subsampling of RNA-Seq read for figure 5 and 6

Subsampling of RNA-Seq reads was conducted with ‘seqtk’ (https://github.com/lh3/seqtk) to normalize total read. The seed parameter, which is specified as -s, was differently specified in each subsampling.

### Calculation and normalization of RNA-Seq count data for figure 3, 4, 5 and 6

All obtained reads were processed with Trimmomatic (version 0.3.3) (Bolger, Lohse and Usadel, 2014) using the following parameters: TOPHRED33 ILLUMINACLIP: TruSeq3-SE.fa:2:30:10 LEADING:19 TRAILING:19 SLIDINGWINDOW:30:20 AVGQUAL:20 MINLEN:40. This procedure removed adapter sequences (ILLUMINACLIP:TruSeq3-PE.fa:2:30:10), leading and trailing low quality or N bases (below quality Phred score 19) (LEADING:19 TRAILING:19). Also, this trimmed the reads when the average quality per base drops below 20 with a 30-base wide sliding window (SLIDINGWINDOW:30:20). Finally, trimmed reads with length > 39 nucleotides and average quality score > 19 were output. The trimmed reads were then mapped to the *O. sativa* reference sequences of IRGSP-1.0_transcript^43, 44^, rRNAs, and transcripts coded in the mitochondria and chloroplast genomes (GenBank; NC_001320.1 and NC_011033.1), the *A. thaliana* reference sequence of Araport11 representative transcripts^45^, the *S. cerevisiae* cDNA R64-1-1 reference sequence of ensembl^46^ or the *D. rerio* GRCz11.cdna.all reference sequence of ensembl^46^ with RSEM (version 1.3.0)^47^, using Bowtie (version 1.1.2)^48^ with default parameters.

In the case of using UMI in the analysis, all obtained paired reads were processed by using dynacomkobe/biodocker_rnaseq_pipeline:ver.0.2.0 with the default parameters, with which quality trimming with Trimmomatic, quantification with RSEM and bowtie were conducted.

Expected counts of each gene in the RSEM outputs were used with R (version 3.4.2)^49^ in the following analysis. The R script used in this study is deposited in xxxxxxx. Briefly, normalization of read counts were conducted by calculating read per million (RPM) by using total read count of all transcripts as denominators. Differentially quantified genes (FDR = 0.05) were detected as described by Sun et al^50^. (2947/3000 words)

## Data availability

All RNA-Seq sequence data is deposited in PRJNA643885 of SRA.

## Acknowledgements

We thank Yoichi Hashida, Yuko Kurita, Yuma Wakamatsu, and Takahiro Mochizuki and Fumiyoshi Abe for their advices on of *A. thaliana* and *O. sativa*, breeding of *D. rerio* and a kind gift of *S. cerevisiae* respectively. We also thank all our laboratory members for their help and encouragement.

---

**Supplemental figure 1.**
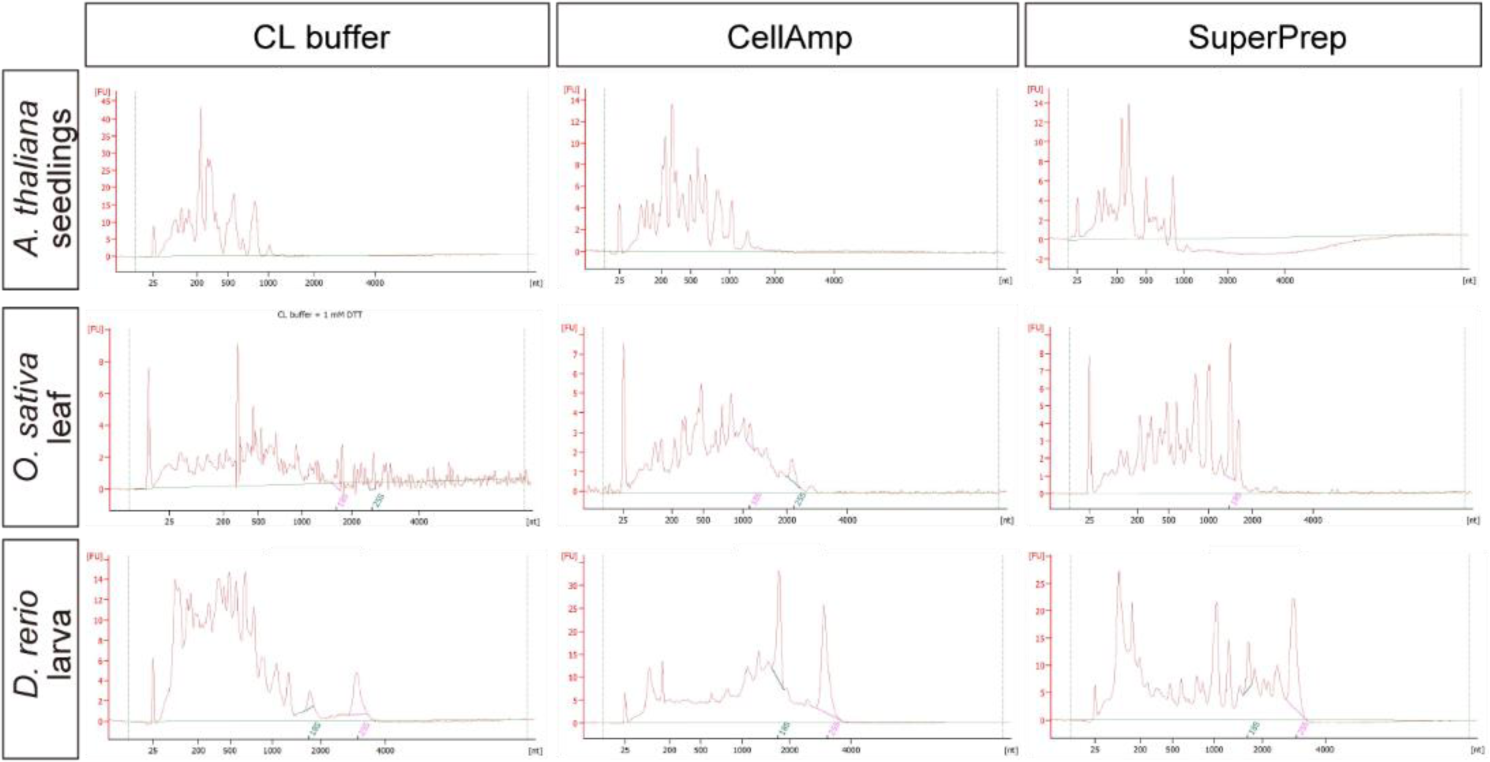
The performance of previously reported buffers for mammalian cultured cells in direct-lysate reverse transcription of *A. thaliana*, *O. sativa* and *D. rerio*. Bioanalyzer electropherograms of purified RNA from plants’ and zebrafish larvas’ lysates homogenized in CL buffer and the buffers in CellAmp kit and in SuperPrep kit.

**Supplemental figure 2.**
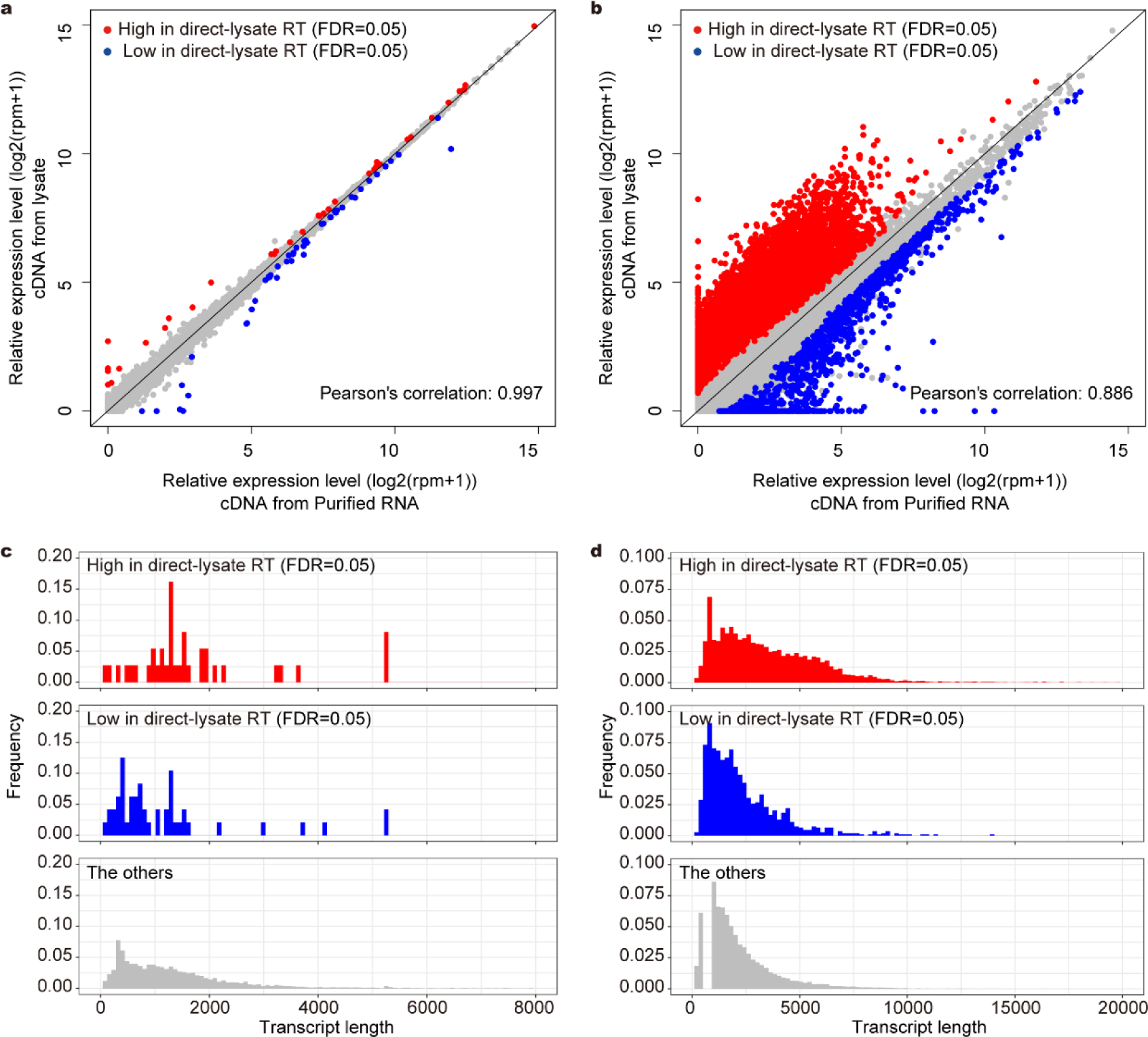
Comparison of RNA-Seq results of purified RNA and lysate of *S. cerevisiae* and *D. rerio*. The results of 3’ RNA-Seq of purified RNA and lysate of *S. cerevisiae* (a, c) and *D. rerio* (b, d) larva. (a, b) Scatter plot of log2 RPM+1 of each gene. Differentially quantified genes (DQGs) were detected between RNA-Seq for purified RNA and lysate (FDR = 0.05, n=6). (c, d) Transcript length distribution of DQGs with higher rpm values in the method using lysate and the other genes. The histograms from 0 nt to 8000 nt in (c) and from 0 nt to 20000 nt in (d) and are shown.

**Supplemental figure 3.**
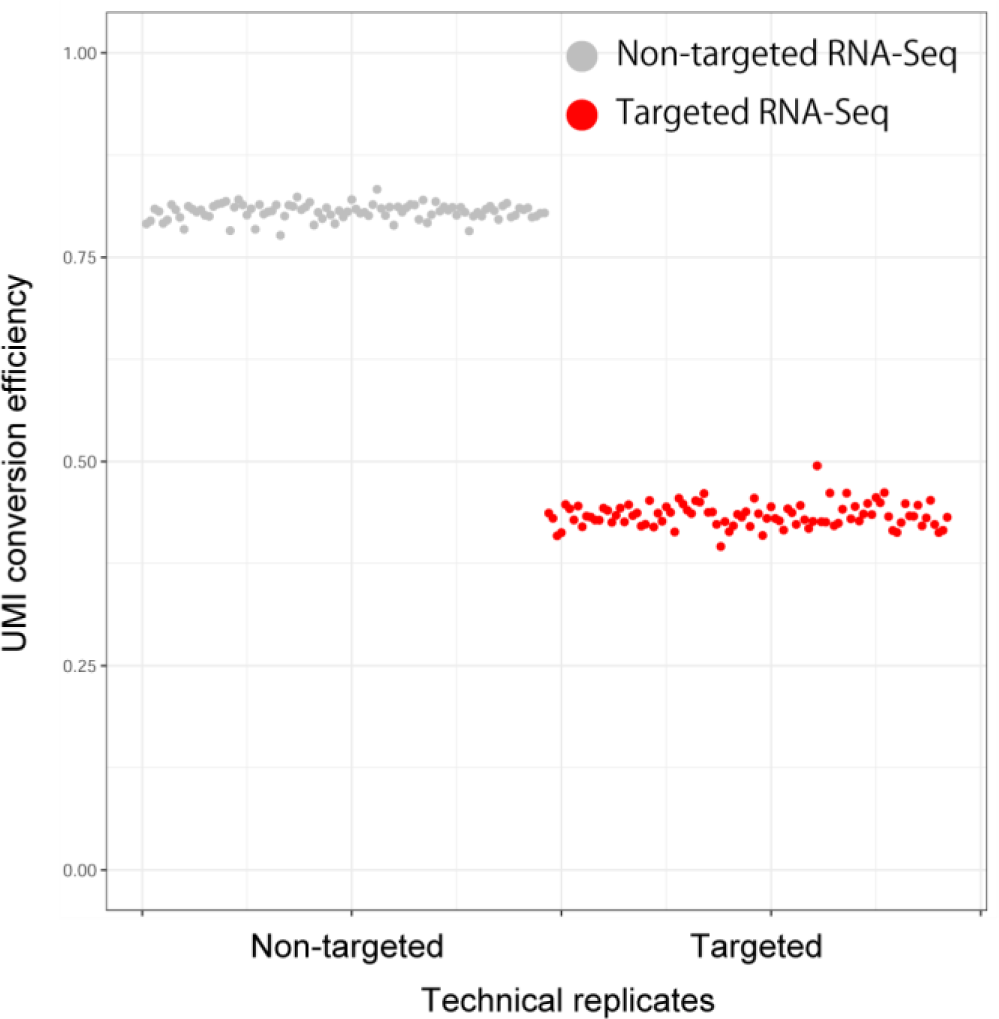
UMI conversion efficiencies of 96 technical replicates of non-targeted and targeted RNA-Seq. A plot of UMI conversion efficiencies of each technical replicates of non-targeted and targeted RNA-Seq in Fig.4. Red indicates targeted RNA-Seq. Gray indicates non-targeted RNA-Seq.

